# Endosperm turgor pressure both promotes and restricts seed growth and size

**DOI:** 10.1101/2021.03.22.436392

**Authors:** Audrey Creff, Olivier Ali, Vincent Bayle, Gwyneth Ingram, Benoit Landrein

**Affiliations:** Laboratoire Reproduction et Développement des Plantes, Univ Lyon, ENS de Lyon, UCB Lyon 1, CNRS, INRA, F-69342, Lyon, France 69364 LYON Cedex 07, France

## Abstract

Organ size depends on complex biochemical and mechanical interactions between cells and tissues. Here, we investigate the control of seed size, a key agronomic trait, by mechanical interactions between two compartments: the endosperm and the testa. By combining experiments with computational modelling, we tested an incoherent mechanical feedforward loop hypothesis in which pressure-induced stresses play two antagonistic roles; directly driving seed growth, but indirectly inhibiting it through mechanosensitive stiffening of the seed coat. We show that our model can recapitulate wild type growth patterns and explain the small seed phenotype of the *haiku2* mutant. Our work further reveals that the developmental regulation of endosperm pressure is needed to prevent a precocious reduction of seed growth rate induced by force-dependent seed coat stiffening.

## Introduction

How tissue growth arrest is achieved once an organ has reached a defined size is a key, yet unresolved, question in developmental Biology (*1*, *2*). In *Drosophila*, mechanical and biochemical signals have been proposed to act in concert to control growth and determine organ size in the wing imaginal disk (*3*, *4*). In plants, mechanical signals can affect growth by modulating key processes such as cytoskeleton organization (*5*, *6*), auxin distribution (*7*, *8*), chromatin organization (*9*) and gene expression (*10*, *11*). However, it remains unclear whether mechanical signals are involved in organ size control in plants.

Seed size is a key agronomic trait that influences seed composition, and viability (*12*). Seed growth relies on interactions between two seed compartments: the endosperm and the testa (*13*). During early post-fertilization development in *Arabidopsis,* the endosperm comprises a single poly-nucleate cell filling most of the internal compartment of the seed (**Fig. 1A**) (*14*). Hydrostatic pressure (turgor) in the endosperm, resulting from osmolite accumulation, is thought to drive seed growth (*15*), while progressive reduction of endosperm turgor was proposed to contribute to seed growth arrest (*15*). However, turgor does not always correlate positively with growth as recently shown in meristematic cells (*16*).

**Fig.1.**
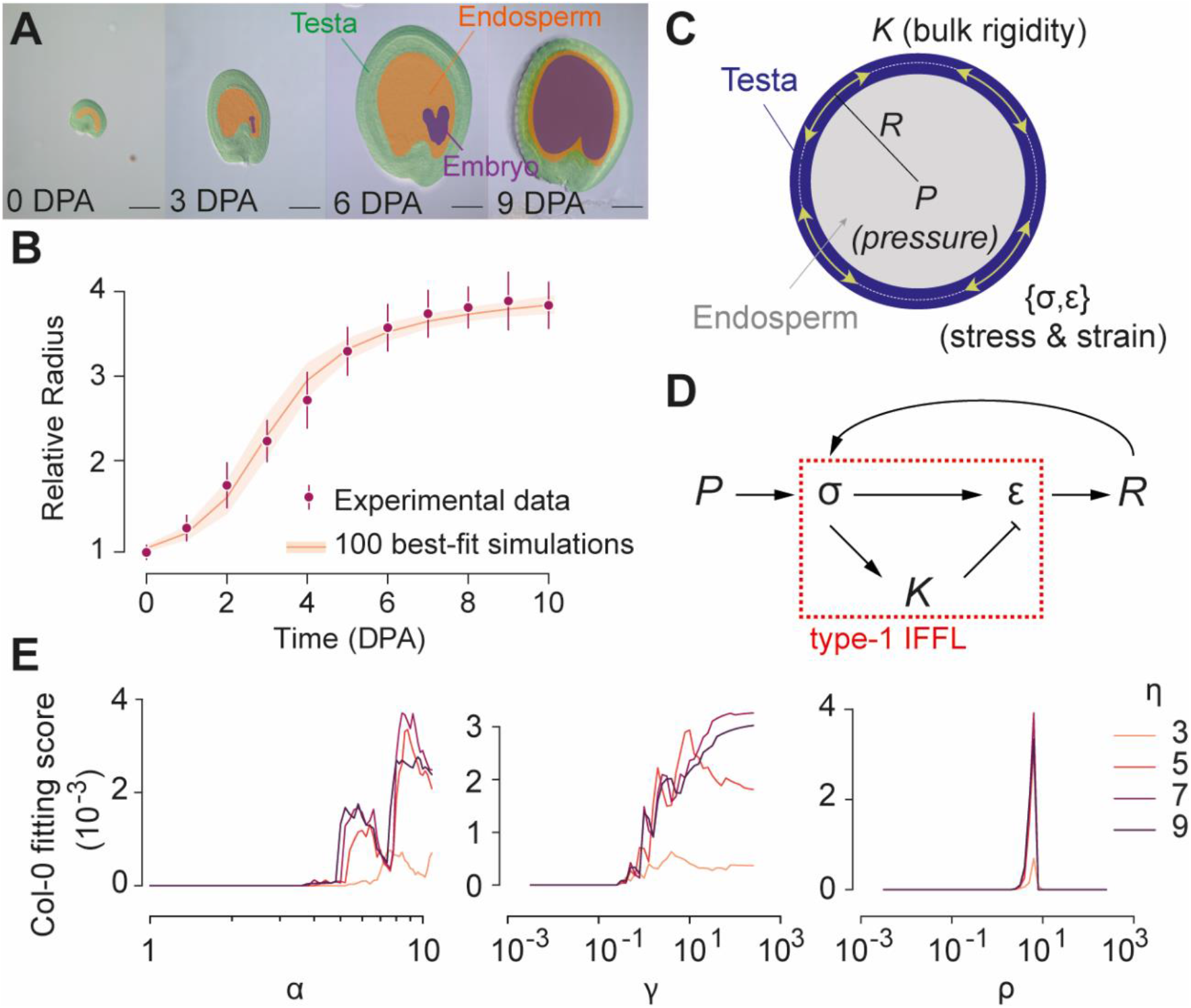
WT seed growth can be modelled using a mechanosensitive incoherent feedforward loop. (**A)** WT seed growth (DPA: Days post-anthesis). Scale bars, 100 μm. (**B**) Relative seed radius as a function of time (DPA). Purple dots and bars correspond to mean and the standard deviation of experimental data for WT seeds (Col-0, pool of 5 independent experiments, 4914 seeds total, 338-464 seeds per day). Orange line and band depict dynamics of 100 simulations that fit best the experimental data. (**C, D**) Modelling seed growth with an incoherent feedforward loop (IFFL). (**E**) Simulation fitting score as a function of four parameters characterizing stress-dependent shell stiffening: (*α*) amplitude of stiffening, (*γ*) threshold ratio between stiffening and growth, (*ρ*) steepness of stiffening mechanics (Hill function exponent), (*η*) characteristic time ratio between growth and stiffening.

The testa, a maternal tissue derived from the ovule chalaza and integuments (*17*), is thought to constrain seed growth. During mid to late seed expansion, the adaxial epidermis of the outer-integument (ad-oi) appears to restrict growth by reinforcing its inward-facing cell wall (wall 3, the third periclinal wall counting from the outside, **Fig. S1**). This process could involve perception of tensile stresses induced in the testa by endosperm pressure. Indeed, the expression of *ELA1 (EUI-LIKE P450 A1*), a negative seed size regulator expressed predominantly in ad-oi (*18*), is promoted by increasing tensile stress in this layer (*11*). Thus seed size could be determined by a mechanosensitive incoherent feedforward loop in which the direct growth-promoting activity of endosperm turgor is antagonized by an indirect growth inhibition resulting from the mechano-sensitive stiffening of testa walls.

## Results and Discussion

To test this possibility, we first analyzed Wild-Type (WT, ecotype Col-0) seed growth at 24h intervals-from anthesis using DIC (Differential Interference Contrast) imaging and seed area measurements (**Fig. 1A, Movie S1**). Seed radius increases about 3.5 times, following a typical and reproducible S-curve, to plateau at around 7 Days Post-Anthesis (DPA) (**Fig. 1B and Fig. S2A**). Seed growth peaked at 1-3 DPA before slowly decreasing between 3 and 7 DPA (**Fig. S2B**). At the end of the growth phase, the endosperm cellularizes, a progressive process which is necessary for subsequent embryo development (*19*). Cellularization has been proposed to influence seed growth arrest because its onset correlates with the end of the growth phase in WT and in some seed size mutants (*13*, *20*), However, we found that seed growth arrest is gradual and starts at least 2 days before the onset of cellularization, which occurs around 5 DPA (**Fig. S2B and Fig. S3A**). We analyzed *ede1-3 (endosperm defective* 1), a mutant lacking a microtubule binding protein required for endosperm cellularization (*19*, *21*). *Ede1-3* mutants only showed a minor seed growth defect (**Fig. S3B-D**), suggesting that endosperm cellularization and seed growth arrest can be dissociated.

Using concepts developed for modelling gene regulatory networks (*22*), we next tested whether seed growth can be formalized through a mechano-sensitive incoherent feedforward loop where endosperm pressure directly sustains seed growth but indirectly inhibits it through force-dependent testa stiffening. We developed a quasi-static morphomechanical model of the seed where the testa is assimilated to a linear elastic spherical shell loaded with pressure forces generated by the endosperm (**Fig. 1C-D**). At mechanical equilibrium, the resulting strain (ε) and stress (σ) induce respectively cell expansion and wall stiffening within the testa. We formalized cell expansion and wall stiffening through a dimensionless system of two coupled ordinary differential equations (see Supplementary Materials). We performed a parameter space exploration on this differential system to identify parameter values yielding simulations compatible with experimental measurements. Among the 5 x 10^5^ parameter sets we tested, less than 2 x 10^3^ yielded simulations converging toward biologically relevant solutions (i.e. R_10DPA_/R_0DPA_ = 3.5 ± 0.5). We quantitatively compared each of these 2 x 10^3^ simulations with experimental data and scored their fit, keeping only the 100 best-fitting simulations (**Fig. 1B**). Analysis of the relationship between fitting score and parameter distribution revealed that the stress-sensitive stiffening would have to be highly non-linear (i.e. strong and sharp) and occur late compared to growth, to account for seed growth control through a mechano-sensitive regulation (**Fig. 1E**). Fitting our simulations to *ede1-3* seed growth dynamics retrieved similar parameter value distributions to those of the WT fit, underlining the similarity between WT and *ede1-3* seed growth (**Fig. S4**).

We next asked whether our model could be used to explain the phenotype of known seed size mutants. We analyzed a mutant allele of *HAIKU2 (IKU2*) (*25*), which encodes a receptor-like kinase only expressed in the endosperm (**Fig. S5A**), and acting in a zygotic growth-control pathway (*15*). Seed growth was initially higher in *iku2* seeds than in the WT, but decreased faster, ultimately leading to the production of smaller seeds (**Fig. 2A and B, Fig S5B**). We hypothesized that *iku2* seed growth defects might result from reduced endosperm turgor (*14*). Using a published method to extract endosperm turgor from force-displacement curves obtained by microindentation (*15*), we confirmed that endosperm pressure decreases throughout the growth phase in WT seeds (**Fig. 2C and Fig. S5C**). To our surprise, this decrease was not observed in the *iku2* mutant, where endosperm pressure was constant and higher than in the WT from the globular stage onward (3 to 4 DPA).

**Fig.2.**
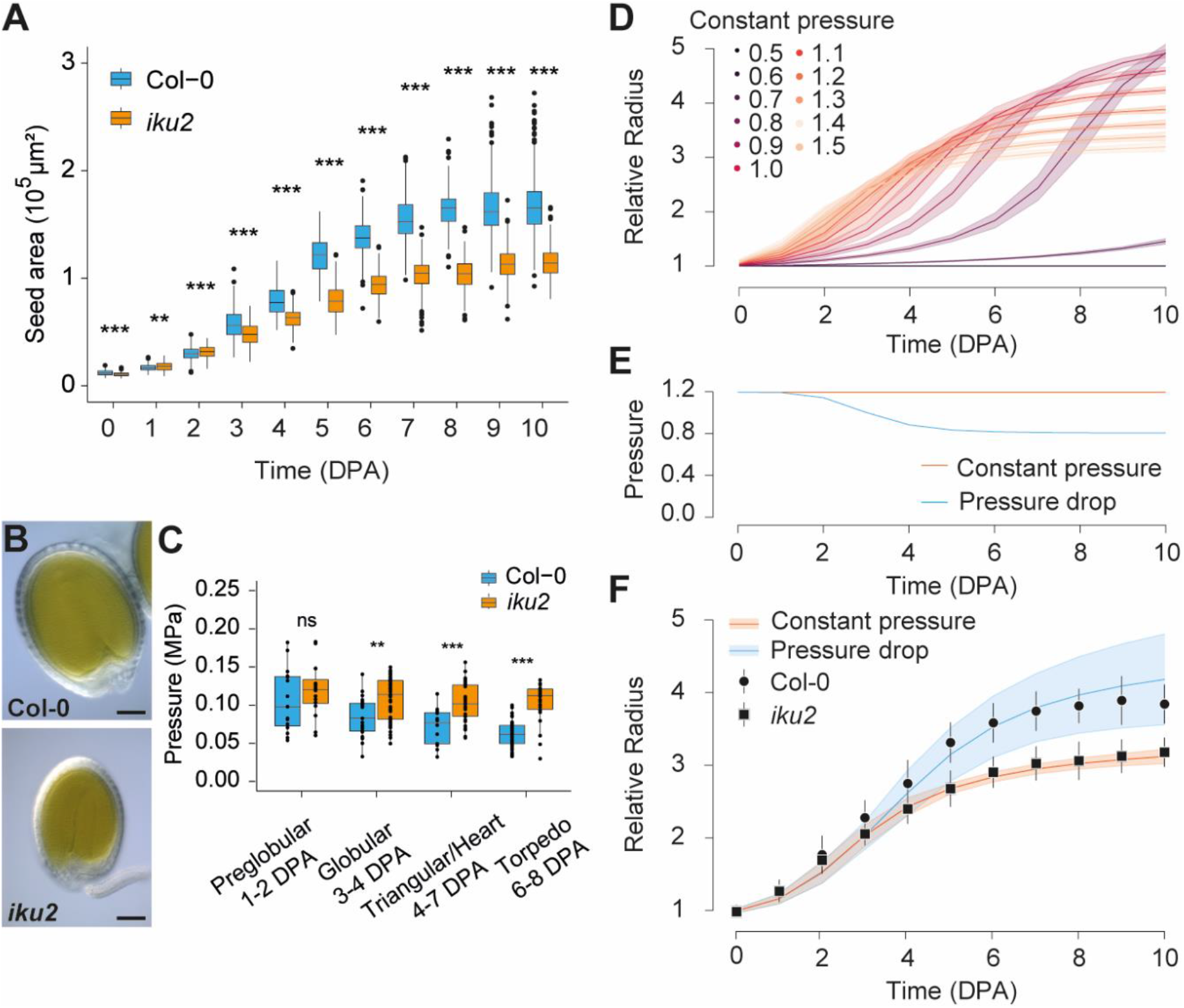
Increased endosperm pressure leads to reduced seed growth in *iku2* mutant. (**A**) WT and *iku2* seed growth (DPA: Days Post Anthesis). Three independent experiments were pooled (Col-0: 2757 seeds total, 216-282 seeds per day, *iku2*: 2783 seeds total, 195-289 seeds per day). Areas were compared using bilateral Student tests. (**B**) Col-0 and *iku2* seeds at 10 DPA. Scale bars: 100 μm. (**C**) Col-0 and *iku2* endosperm pressure. (Col-0: 103 seeds total, 14-51 seeds per stage; *iku2*: 126 seeds total, 16-51 seeds per stage). Pressure values were compared using bilateral Student tests. (**D**) Relative radius as a function of time (in DPA) and pressure in simulations. Thick curves and shadowed bands correspond to the mean behavior and standard deviation of the 100 simulations best-fitting WT experimental data. (**E**) Pressure drop implemented to reproduce pressure changes observed in WT seeds. (**F**) Relative seed radius as a function of time in Col-0 and *iku2*. Black circles and squares with bars represent mean seed size and standard deviation in Col-0 and *iku2* measured experimentally (panel A). Solid lines and shadowed bands show mean and standard deviation of the 100 best simulations where the model was fitted to *iku2* at a constant pressure of 1.2 or where a pressure drop (E) was implemented.

To better understand this counterintuitive result, we tested how endosperm turgor affects growth in simulations. For small values of our dimensionless pressure variable (below 0.7), growth was, as expected, either null or limited (**Fig. 2D, Fig. S6A**). However, for higher values (above 0.7), increasing pressure initially induces faster initial growth but leads to smaller final radii. We then tested whether the sustained endosperm pressure observed in *iku2* could explain the *iku2* growth phenotype. We fitted the model at constant pressure to the experimental measurements of *iku2* seed growth, which did not alter the global behavior of the system (**Table S4, Fig. S7)**. We then applied a step function to mimic the pressure reduction measured experimentally in WT seeds to these *iku2* fitted simulations (**Fig.2E**). This led to an extension of the growth phase in the *iku2* fitted simulations which, strikingly, could now recapitulate experimentally measured WT growth patterns (**Fig. 2F**).

To test if turgor-induced changes in growth could be linked to stress-dependent stiffening of the testa, we first tested the influence of pressure on shell stiffness in simulations. Increased pressure led to precocious shell stiffening (**Fig.3A, Fig. S6B**), explaining the previously observed growth reduction (**Fig. 2D, Fig. S6A**). In contrast, decreasing pressure over time delayed shell stiffening (**Fig.3B**). We then tested experimentally whether *iku2* seed growth defects could be due to precocious stress-dependent testa stiffening. We previously showed that the expression of the mechanosensitive gene *ELA1* is increased in the *iku2* mutant by qPCR (*11*). To confirm this, we quantified *pELA1::3X-VENUS-N7* reporter fluorescence in WT and *iku2* seeds. We observed higher fluorescence in the oi-ad layer of *iku2* mutant seeds than in WT seeds at all relevant stages of development, suggesting that increased endosperm pressure in *iku2* increases mechanical response in the testa (**Fig.3C-D, Fig. S8**). We then addressed possible alterations of the mechanical properties of testa walls in *iku2* seeds. The pectin matrix is thought to be a key determinant of cell wall mechanical properties. Homogalacturonans (HG), the most abundant pectins, are deposited in a methyl-esterified state, but can subsequently be demethyl-esterified (*23*). This process can promote enzymatic HG degradation (*23*), weakening the cell wall and promoting growth (*24*). However, fully demethylesterified HGs can form calcium-dependent cross-links that increase wall stiffness and inhibit growth (*25*). Using three different antibodies (LM19, JIM5 and 2F4), we assessed HG methylesterification in seeds at different growth stages by immunolocalization and subsequent signal quantification in the outer periclinal walls of the testa using a custom-made pipeline (**Fig. S9**). JIM5 preferentially detects pectins with low levels of methyl-esterification (*26*) while LM19 and 2F4 preferentially detect demethylesterified pectins (*27*, *28*). In WT seeds, epitopes for all three antibodies were more abundant in wall 3 than to other walls, supporting its load-bearing role (*11*) (**Fig. 3E to H, Fig S10-12**). At 3 and 4 DPA, all signals were weak and spotty (especially for JIM5 and 2F4), but strongly increased between 4 and 6 DPA, consistent with model predictions that testa wall stiffening occurs late compared to growth, and should be strong and sharp (**Fig 3B**). For all antibodies, the signal was stronger in *iku2* than in Col-0 at early stages of development (from 3 to 5 DPA depending on the antibody) but was similar at the end of the growth phase for JIM5 and 2F4 (between 7 and 9 DPA). *iku2* seed growth restriction could therefore involve precocious pectin demethylesterification and associated stiffening (**Fig. 3I**).

**Fig.3.**
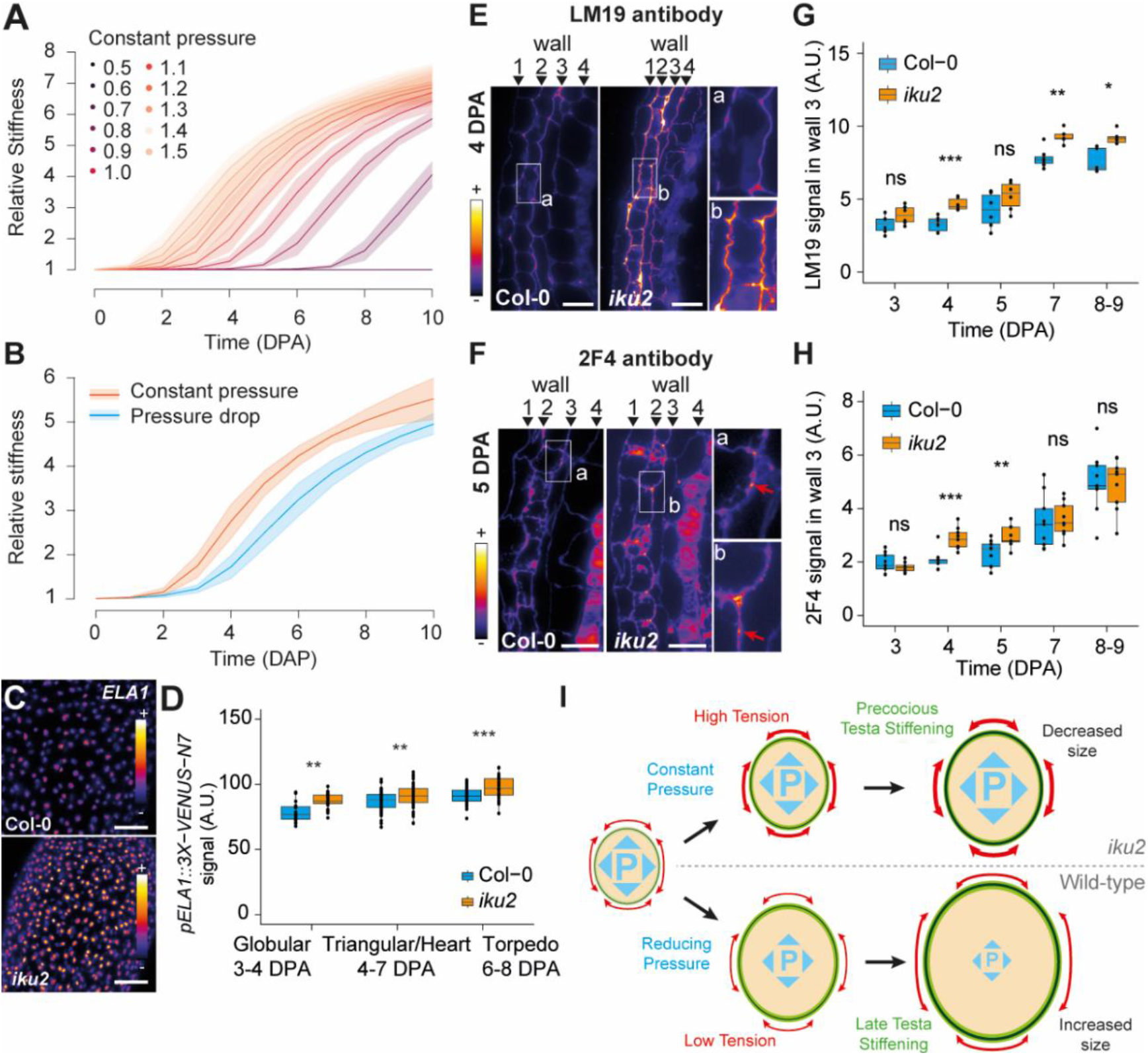
Increased endosperm turgor leads to precocious testa stiffening in *iku2*. (**A**) Relative stiffness (in DPA: Days after pollination) and pressure in simulations. Thick curves and shadowed bands correspond to mean behavior and standard deviation of 100 simulations best-fitting the WT experimental data. (**B**) Relative stiffness as a function of time in 100 best simulations after model fitting to *iku2* data at constant pressure of 1.2 or with pressure drop (Fig. 2E) implementation. (**C**) *pELA1::3X-VENUS-N7* expression in Col-0 and *iku2* seeds at heart embryo stage. Scale bars: 50 μm. (**D**) Mean signal of *pELA1::3X-VENUS-N7* reporter in nuclei of Col-0 and *iku2* seeds (intensity unit/pixel) in two independent experiments. Seeds were classified according to embryo developmental stage (B. Col-0: 24199 nuclei from 129 seeds total, 15-68 seeds per stage; *iku2*: 26623 nuclei from 113 seeds total, 25-58 seeds per stage). Fluorescent signals were compared using bilateral Student tests. (**E and G**) Signal from LM19 (**E**) and 2F4 (**G**) immunolocalizations on Col-0 and *iku2* testas at 4 DPA and 5 DPA respectively. Red arrows show spotty 2F4 signal. Scale bars: 20 μm. (**F and H**) Signal intensity from LM19 (**F**) and 2F4 (**H**) immunolocalizations (x 10^3^ intensity unit/pixel) in wall 3 of Col-0 and *iku2* seeds as a function of time (LM19: pool of two independent experiments, Col-0: 30 seeds total, 5-7 seeds per day; *iku2*: 29 seeds total, 5-6 seeds per day. 2F4: pool of three independent experiments, Col-0: 45 seeds total, 9 seeds per day; *iku2*: 45 seeds total, 9 seeds per day). Signal intensities were compared using bilateral Student tests. **(I)** Antagonistic roles of endosperm turgor in the regulation of seed growth.

The counterintuitive consequences of the endosperm pressure drop in WT seeds, mediated by the *IKU* pathway (*12*) and exposed by our results, have profound implications for crop improvement strategies, particularly those altering fluxes of osmotically active metabolites to enhance seed development. Beyond seeds, the parsimony of our core assumptions suggests that the mechanosensitive incoherent feed-forward motif could be a ubiquitous regulator of plant organogenesis.

## Acknowledgments

We thank Frederic Berger and Claudia Köhler for providing the *iku2* and ede1*-3* mutant seeds respectively; R. Azais, F. Ingels, Y. Long, A. Boudaoud, C. Godin and O. Hamant for helpful discussions and comments on the manuscript; L. Beauzamy for technical assistance regarding indentation; G. Cerutti for technical assistance regarding the distribution of the code; A. Lacroix, P. Bolland and J. Berger for technical assistance regarding plant cultivation; I. Desbouchages and H. Leyral for technical assistance regarding molecular biology work; C. Vial, L. Grangier, N. Camilleri and S. Maurin for administrative assistance; the PLATIM for technical assistance regarding the microscopy.

## Author contributions

O.A., G.I. and B.L. led the study, obtained funding and supervised the work. A.C and B.L carried the experiments. A.C, V.B. and B.L. analyzed the experiments. O.A. developed the computational model and performed the simulations. O.A., G.I. and B.L. wrote the paper with input from all authors. Authors declare that they have no competing interests.

## Data and material availability

All lines used in this study will be provided upon signature of an appropriate material transfer agreement. All the code is available at https://gitlab.inria.fr/mosaic/publications/seed_sup_mat. All the data is available in the main text, in the supplementary materials or at https://zenodo.org/record/4620948#.YFR0Hi1h0UE.

---

**Fig.S1.**
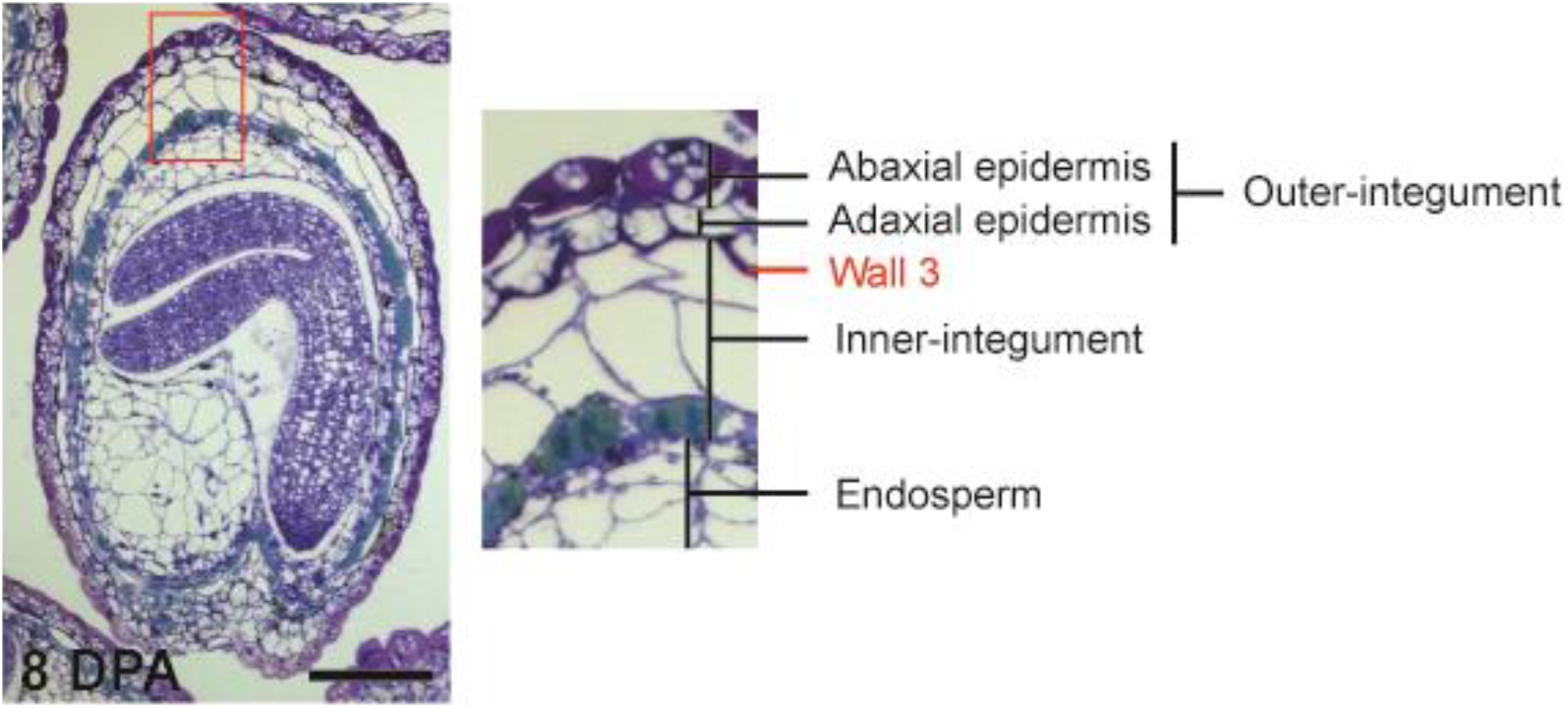
Organization of the testa layers in *Arabidopsis* seeds. Col-0 seed at 8 DPA (Days after pollination) stained with toluidine blue. The close-up view shows the organization of the testa layers and the presence of wall 3 separing the innner and the outer-integuments. Scale bar: 100 μm.

**Fig.S2.**
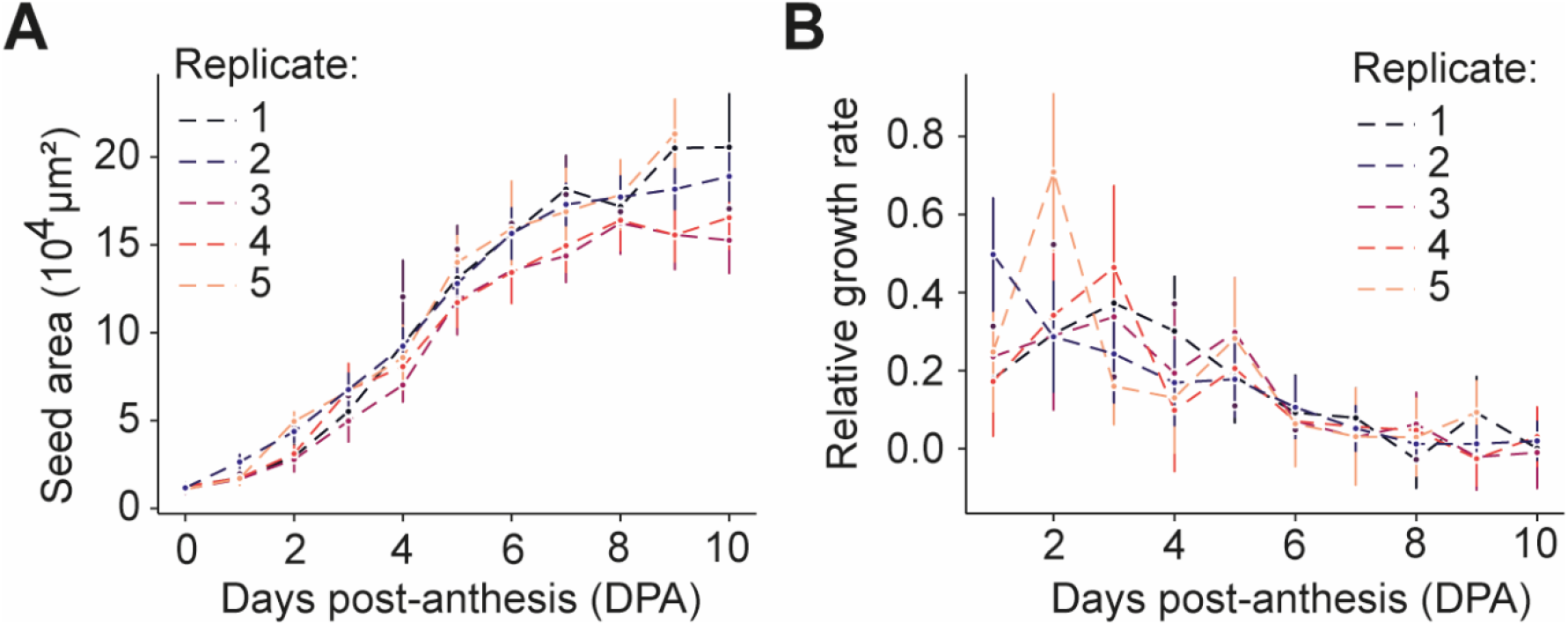
Quantification of WT seed growth pattern. Mean WT seed area (**A**) and relative growth rate (**B**) as a function of time (in DPA) in 5 independent experiments (782-1083 seeds total per experiments, 33-110 seeds per day per experiment). Error bars show the standard deviation.

**Fig. S3.**
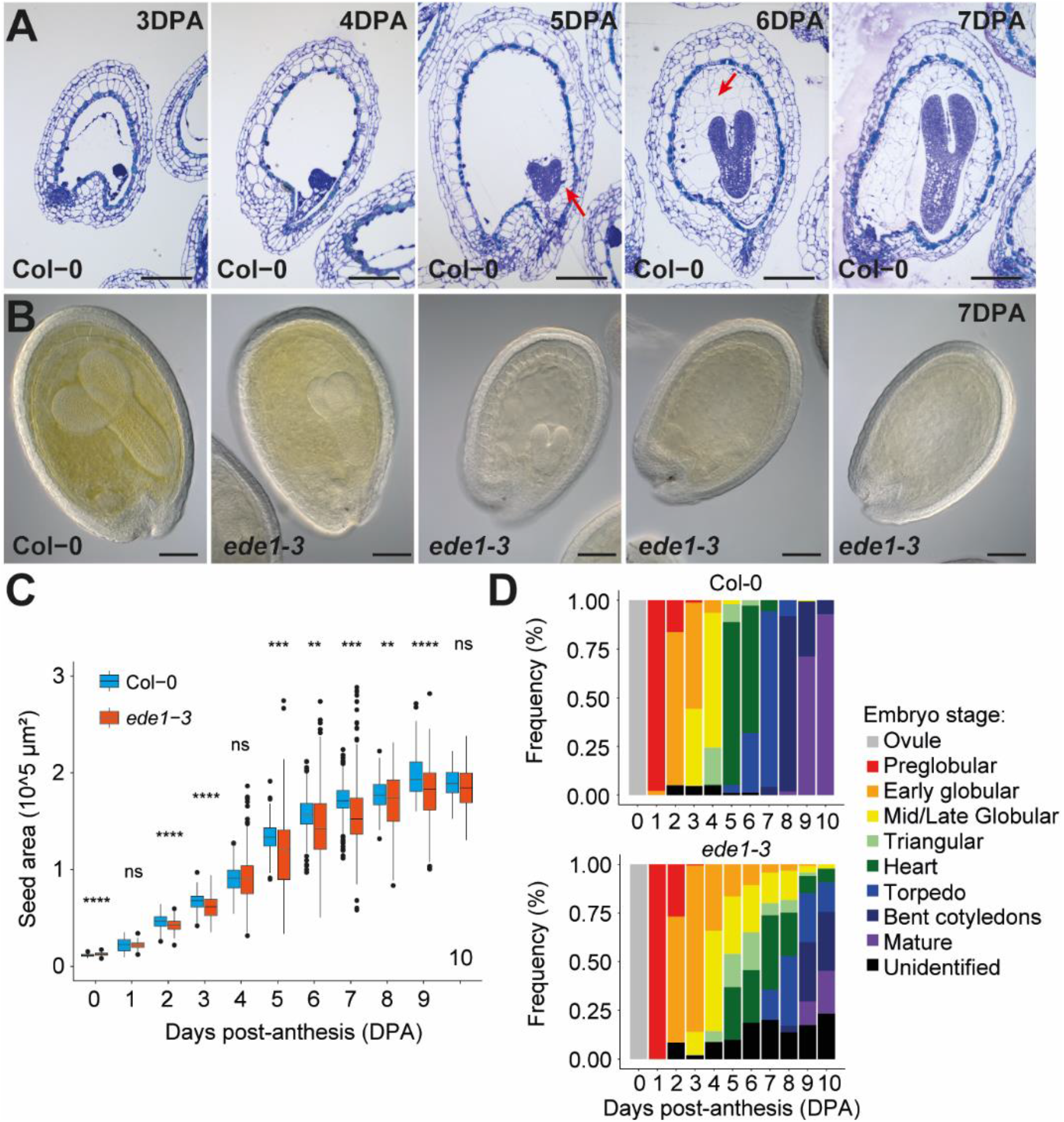
Endosperm cellularization does not control seed growth arrest. (**A**) WT seeds stained with toluidine blue at different stages of development (in DPA: days post-anthesis). The red arrows mark the initiation and progression of cellularization from 5 DPA onwards. Scale bars: 100 μm. (**B**) WT and *ede1-3* at 7 DPA showing examples of defects in embryo growth in the mutant. Scale bars: 100 μm. (**C**) Seed area as a function of time (in DPA) in Col-0 and *ede1-3*. Two independent experiments were pooled (Col-0: 1843 seeds total, 84-195 seeds per day, *ede1-3*: 1584 seeds, 86-187 seeds per day). Error bars show the standard deviation. Seed areas were compared using bilateral Student tests. (**D**) Stage classification of the embryos in the batch of Col-0 and *ede1-3* seeds presented in (C) (Col-0: 1843 seeds, 84-195 seeds per day, *ede1-3*: 1584 seeds, 86-187 seeds per day). The “unidentified” class corresponds to seeds where the embryo was not visible or, in the case of *ede1-3* at late stages of development, displayed defects similar to those presented in panel B that precluded the classification.

**Fig.S4.**
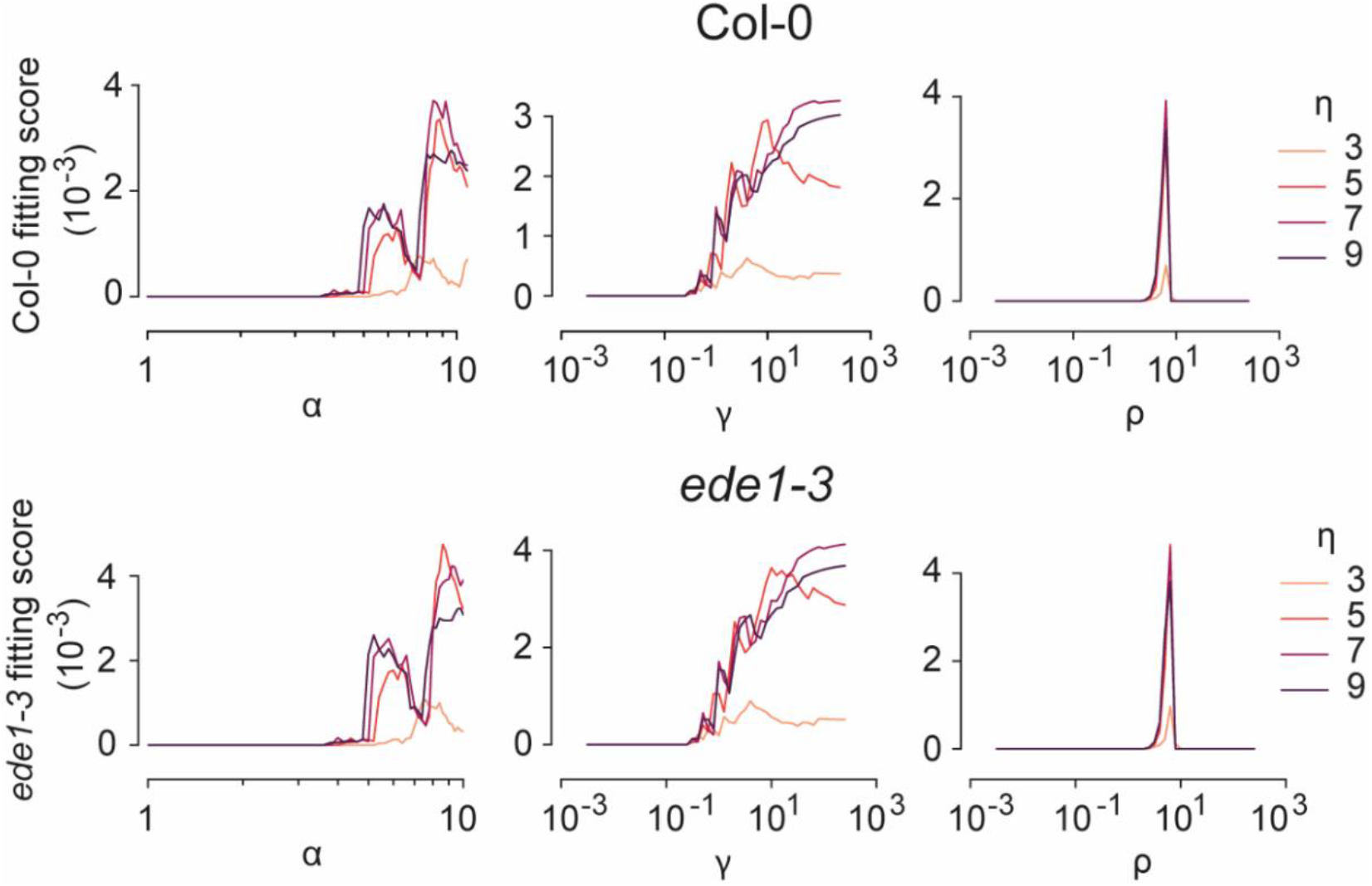
The simulations fit similarly to Col-0 and *ede1-3* data. Fitting score of the simulations to Col-0 and *ede1-3* experimental data as a function of 4 key stress-dependent stiffening parameters of the model. (α) mechano-sensitivity of the stiffening pathway, (γ) threshold ratio between stiffening and growth, (ρ) Hill function exponent, steepness of the mechanosensitive stiffening mechanics, (η) characteristic time ratio between growth and stiffening.

**Fig.S5.**
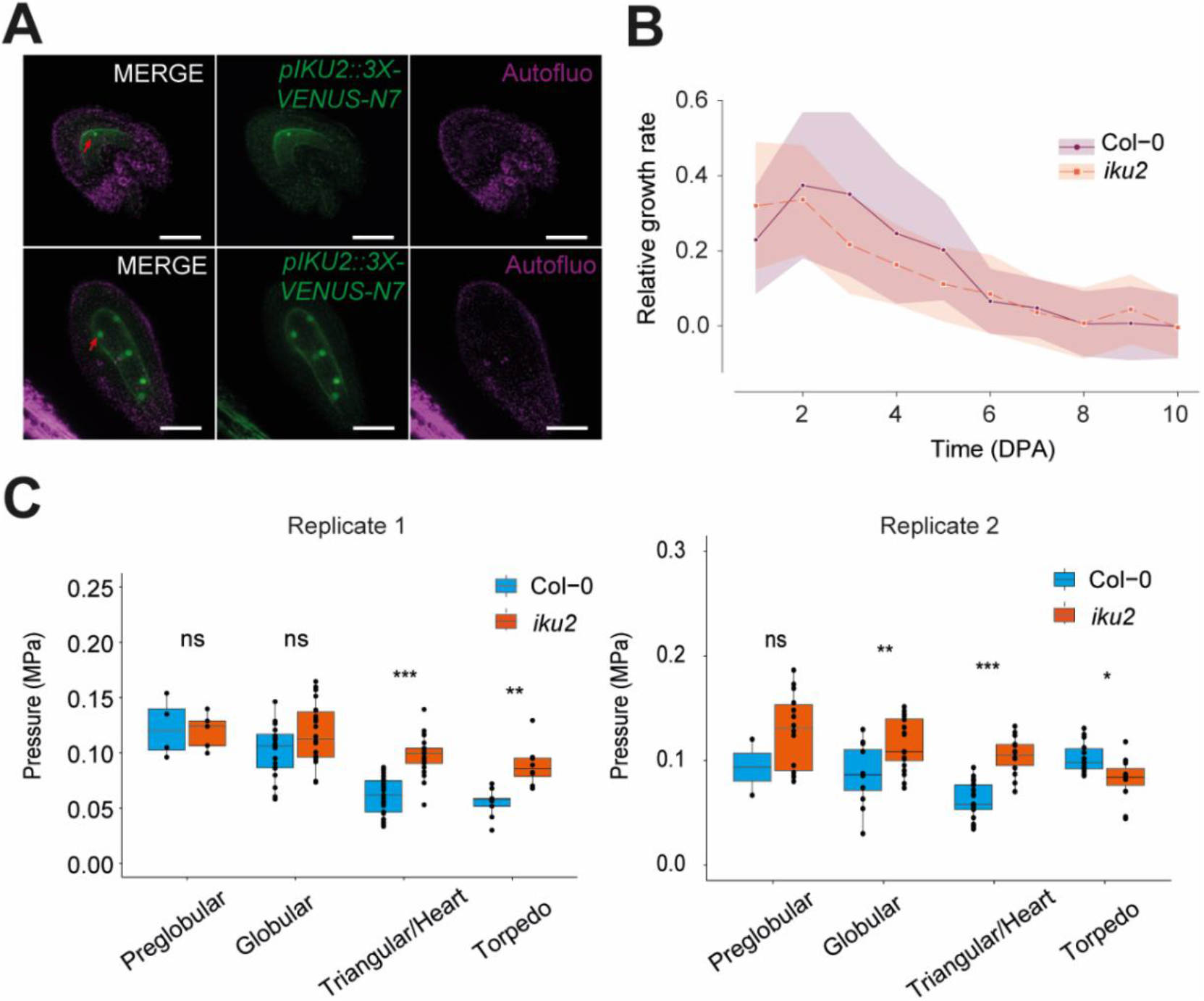
Increased pressure leads to reduced seed growth in *iku2* mutant. (**A**) Signal of the *pIKU2::3X-VENUS-N*7 reporter in seeds at 1 to 2 DPA showing the presence of fluorescent signal in endosperm nuclei only. Scale bars, 50 μm. (**B**) Relative seed growth rate of Col-0 and *iku2* obtained by deriving the seed size measurements presented in Fig.2E. The thick curves correspond to the mean behavior and the dim bands to the calculated deviation (see Material and Methods). Three independent experiments were pooled (Col-0: 2757 seeds, 216-282 seeds per day, *iku2*: 2783 seeds, 195-289 seeds per day). (**C**) Endosperm pressure in Col-0 and *iku2* seeds extracted from stiffness measurements measured using a microindentor in two experiments independent from that presented in Fig.2C. Seeds were classified by the developmental stage of their embryo (Replicate 1: Col-0: 132 seeds total, 10-66 seeds per stage; *iku2*: 97 seeds total, 8-54 seeds per stage, Replicate 2: Col-0: 49 seeds total, 2-21 seeds per stage; *iku2*: 59 seeds total, 11-20 seeds per stage). Pressure values were compared using bilateral Student tests.

**Fig.S6.**
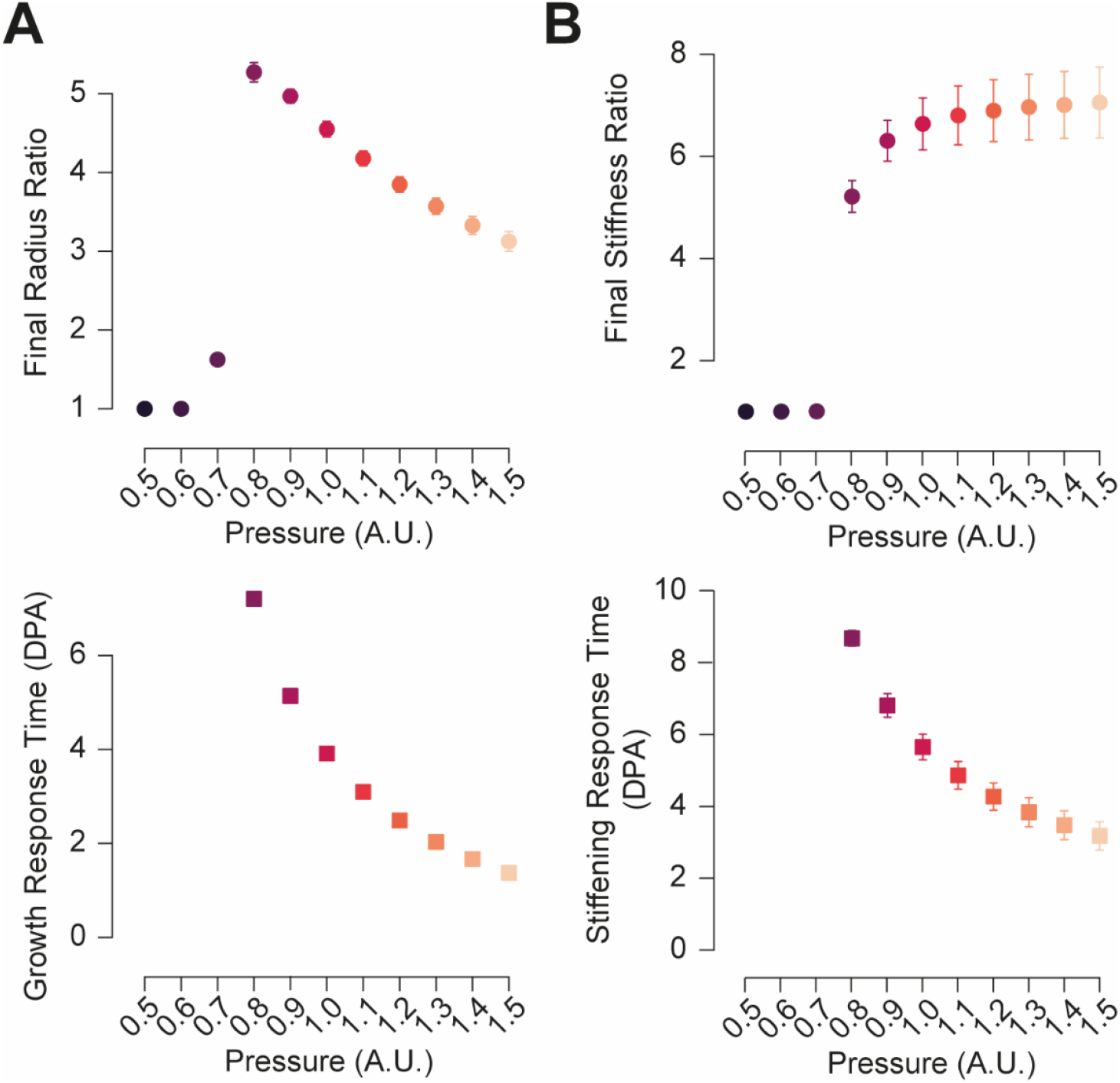
Effect of modulations of pressure on growth and stiffening in simulations. Evolution of radius and stiffness steady state and response time of the expanding shell when loaded with increasing values of pressure. (**A and C**): Ratio between the final and initial values of the relative radius (A) and stiffness (C) for increasing values of pressure. (**B and D**): Response time (defined as the time needed to reach half of the steady state value) of the growth process (B) and the stiffening process (D) for increasing values of pressure. Each point corresponds to 100 simulations, performed with the same constant value of pressure. Error bars depict the standard deviation. In B and D, the first three points of each graph, corresponding to pressure values of 0.5 to 0.7 are missing as the system does not evolve for these small values of pressure.

**Fig.S7.**
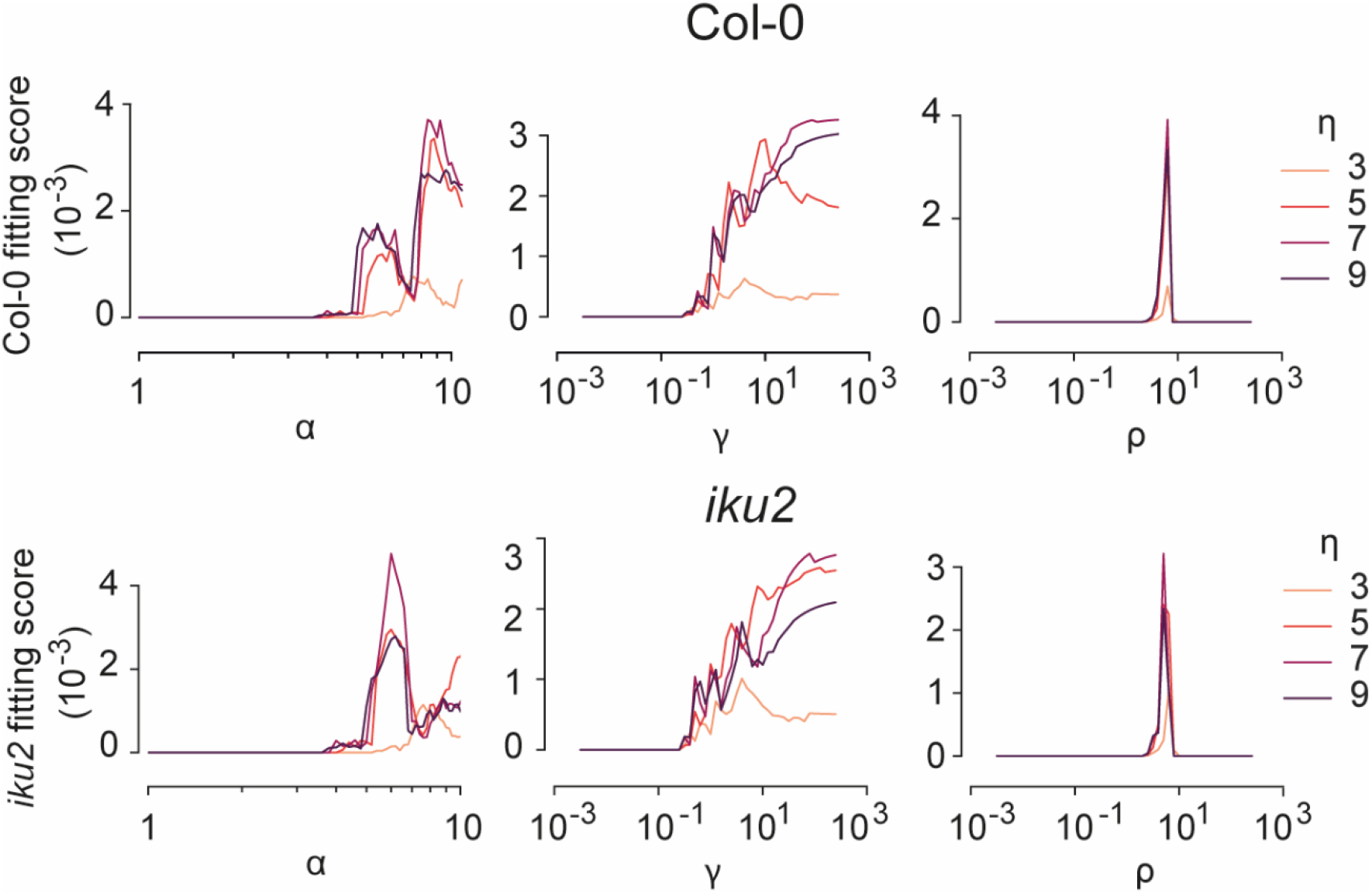
Fitting of the model to *iku2* experimental data. Fitting score of the simulations to Col-0 and *iku2* experimental data as a function of 4 key stress-dependent parameters of the model. (α) mechano-sensitivity of the stiffening pathway, (γ) threshold ratio between stiffening and growth, (ρ) Hill function exponent, steepness of the mechanosensitive stiffening mechanics, (η) characteristic time ratio between growth and stiffening.

**Fig.S8.**
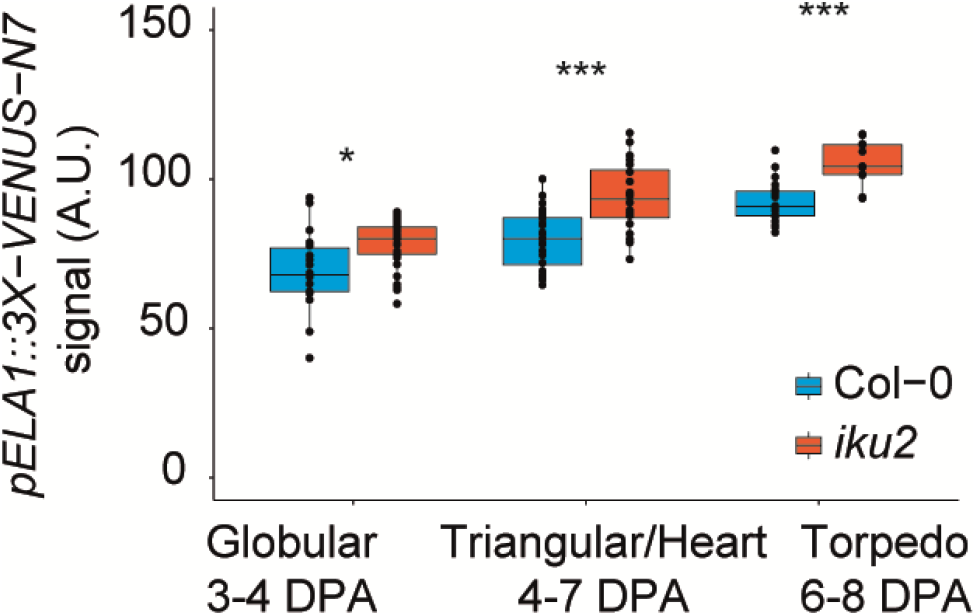
*ELA1* expression is higher in *iku2* seeds. Mean signal of the *pELA1::3X-VENUS-N7* reporter in nuclei of Col-0 and *iku2* seeds (intensity unit/pixel) in an experiment independent from the one presented in Fig.3D. Seeds were classified according to the developmental stage of their embryo (Col-0: 15167 nuclei from 69 seeds total, 18-28 seeds per stage; *iku2*: 18563 nuclei from 67 seeds total, 12-35 seeds per stage). Fluorescent signals were compared using bilateral Student tests.

**Fig. S9.**
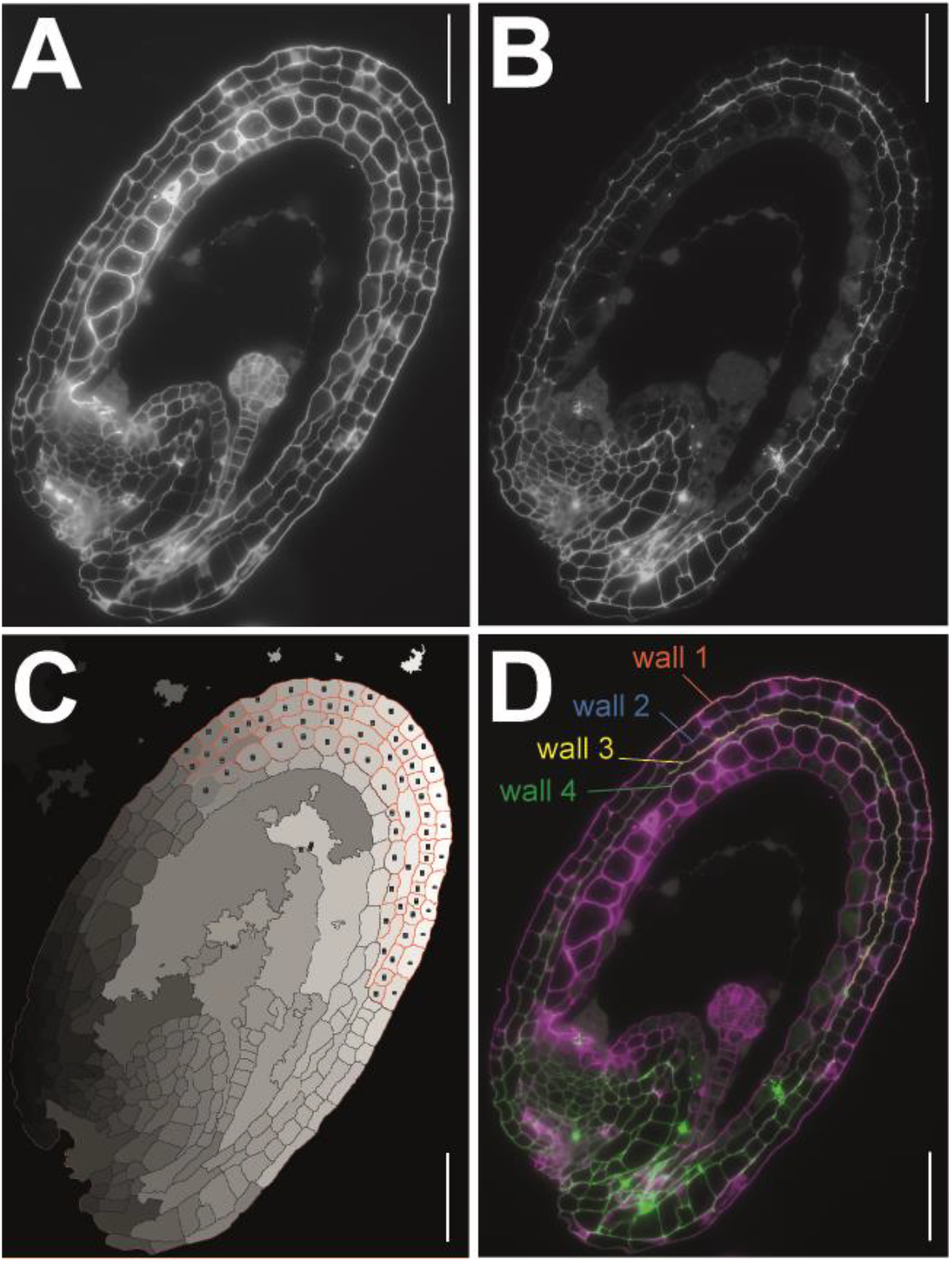
Quantification of immunofluorescence signal in testa walls following immunostaining of cell wall components. (**A**) Control channel (stained with calcofluor). (**B**) Signal channel (stained with the LM19 antibody). (**C**) Segmentation of testa cells using ImageJ and layer assignment. (**D**) Extraction and classification of the periclinal walls of the testa. Final overlay shows the cell wall ROI (Regions Of Interest) used for the quantification. Scale bars, 50 μm.

**Fig.S10.**
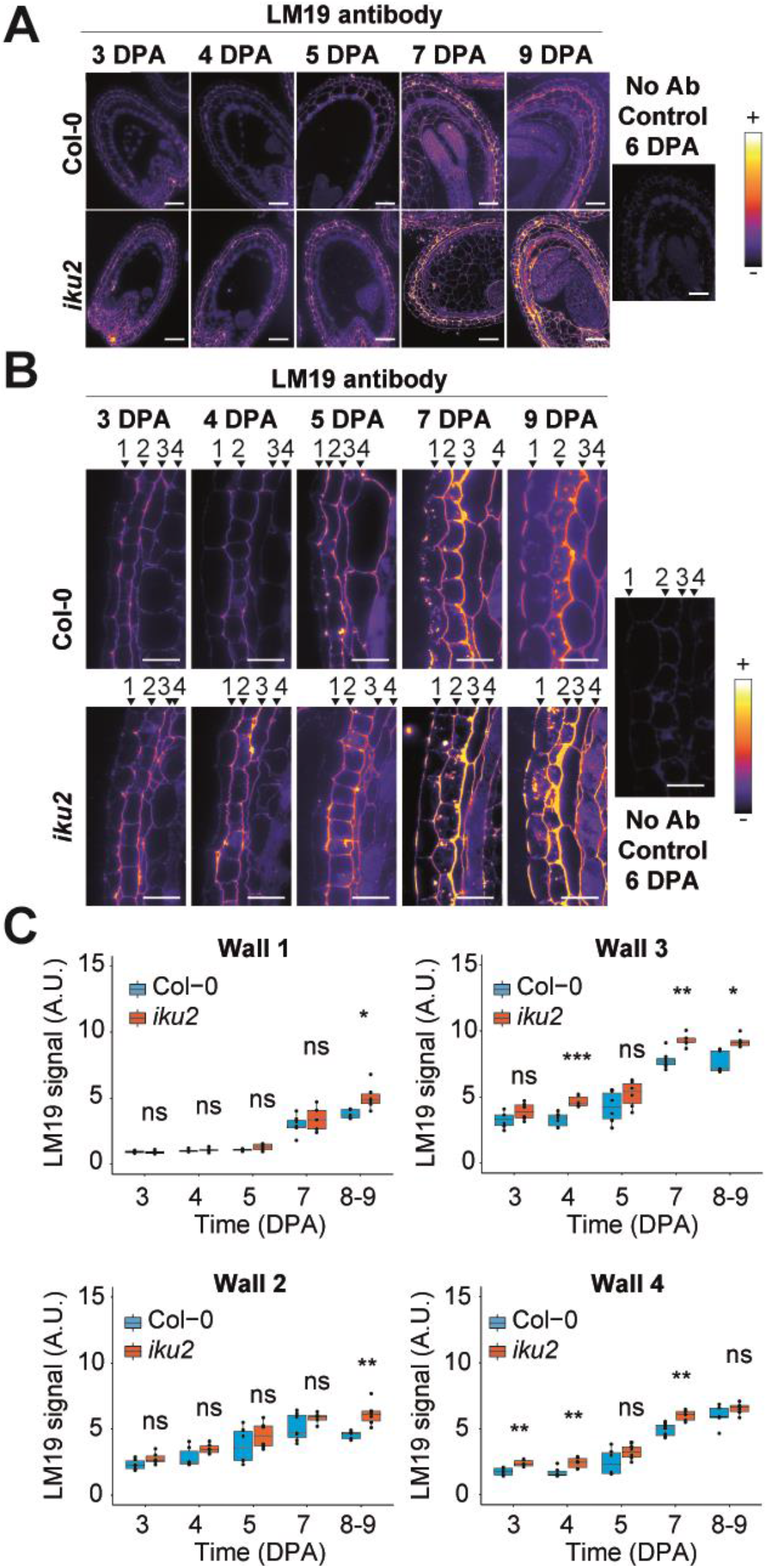
LM19 signal in testa walls. (**A,B**) Signal of the LM19 antibody in Col-0 and *iku2* testa walls obtained by immunolocalization of seed sections at different stages of development (intensities are color-coded using the fire lookup table). Wall numbers (1 to 4) are displayed in the close-up views (B). Scale bars: A. 50 μm and B. 20μm. (**C**) Signal intensity (x 10^3^ intensity unit/pixel) of the LM19 antibody in periclinal testa walls 1 to 4 (counting from the outside of the seed) of Col-0 and *iku2* seeds as a function of time (pool of two independent experiments, Col-0: 30 seeds total, 5-7 seeds per day; *iku2*: 29 seeds total, 5-6 seeds per day). Signal intensities were compared using bilateral Student tests.

**Fig.S11.**
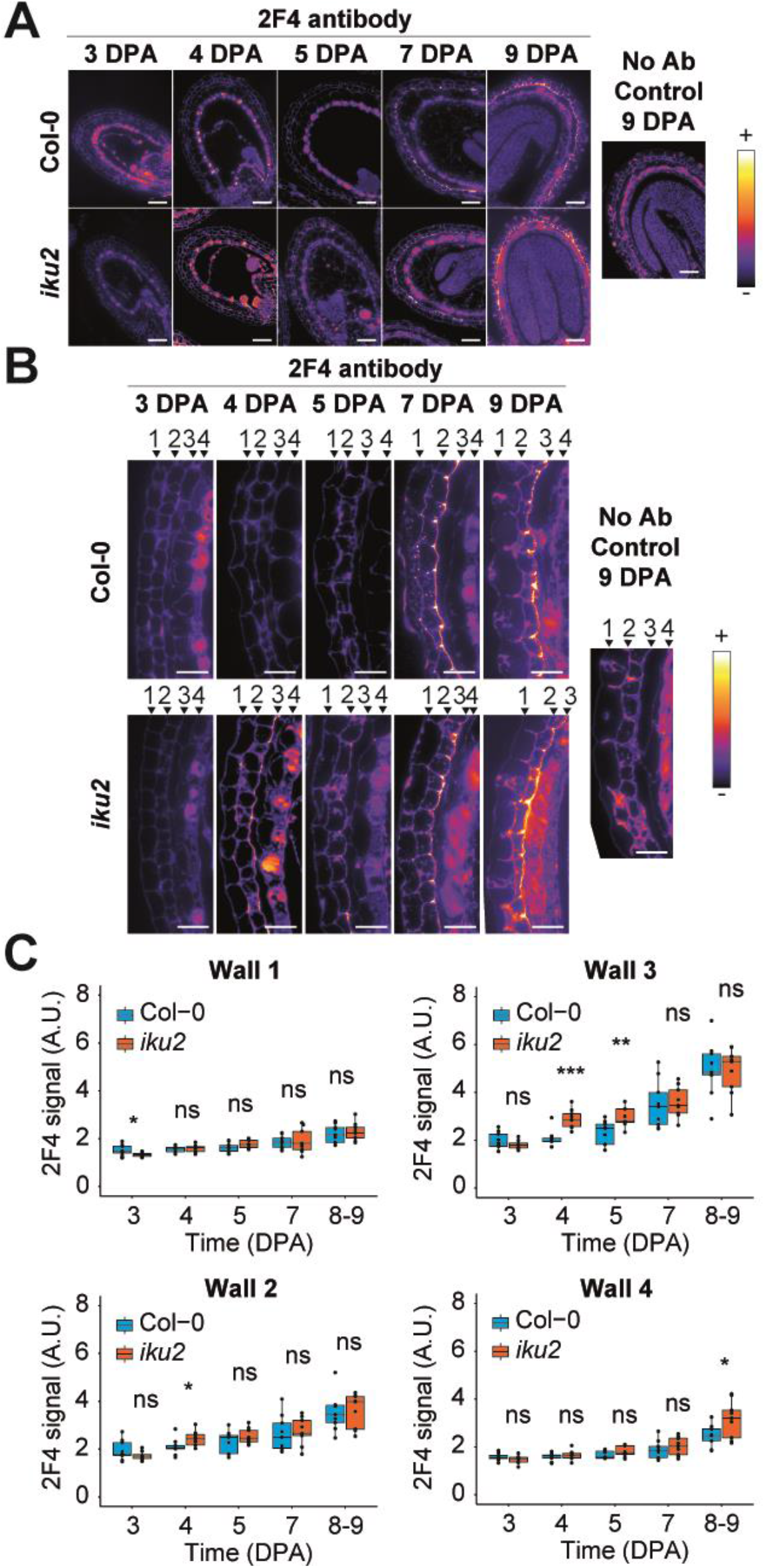
2F4 signal in testa walls. (**A,B**) Signal of the 2F4 antibody in Col-0 and *iku2* testa walls obtained by immunolocalization of seed sections at different stages of development (intensities are color-coded using the fire lookup table). Wall numbers (1 to 4) are displayed in the close-up views (B). Scale bars: A. 50 μm and B. 20μm. (**C**) Signal intensity (x 10^3^ intensity unit/pixel) of the 2F4 antibody in periclinal testa walls 1 to 4 (counting from the outside of the seed) of Col-0 and *iku2* seeds as a function of time (pool of three independent experiments, Col-0: 45 seeds total, 9 seeds per day; *iku2*: 45 seeds total, 9 seeds per day). Signal intensities were compared using bilateral Student tests.

**Fig.S12.**
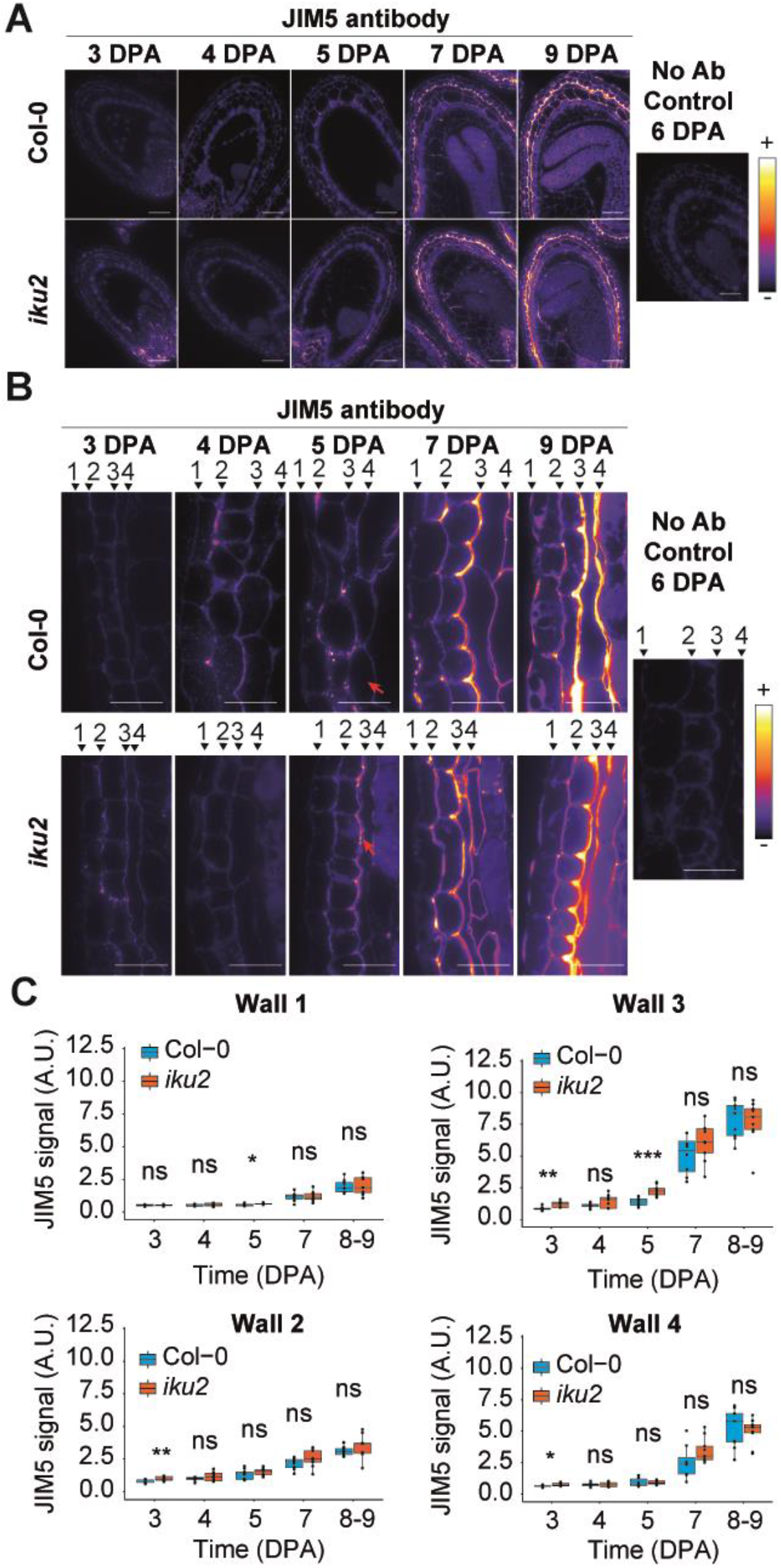
JIM5 signal in testa walls. (**A**) Signal of the JIM5 antibody in Col-0 and *iku2* testa walls obtained by immunolocalization of seed sections at different stages of development (intensities are color-coded using the fire lookup table). Wall numbers (1 to 4) are displayed in the close-up views (B). Scale bars: A. 50μm and B. 20μm. (**C**) Signal intensity (x 10^3^ intensity unit/pixel) of the JIM5 antibody in periclinal testa walls 1 to 4 (counting from the outside of the seed) of Col-0 and *iku2* seeds as a function of time (pool of three independent experiments, Col-0: 45 seeds total, 8-10 seeds per day; *iku2*: 45 seeds total, 8-10 seeds per day). Signal intensities in Col-0 and *iku2* were compared using bilateral Student tests.

## Materials and Methods

### Plant materials and growth conditions

The *iku2-2* and *ede1-3* mutant alleles were described previously (*1*, *2*). The fluorescent marker lines *Lti6b:GFP* and *pELA1::3X-VENUS-N7* were also described previously (*3*, *4*). The *pIKU2::3X-VENUS-N7* was developed for this study (see below).

Seeds were gas sterilized with chlorine (3mL HCl (37%) in 150mL bleach) for 2 hours and sown on plates with Murashige and Skoog (MS) medium and 0,5% sucrose in sterile condition, stratified for 2 days at 4°C and grown for 11 days in a Sanyo (Fisher Scientific) under short-day conditions (8 h light, 21°C during the day, 18°C during the night, 150 μmol.m^−2^.s^−1^). Seedlings were then transferred into separate pots of soil (Argile 10 (Favorit)), and put in a short day growth chamber (8 h light, 21°C during the day, 18°C during the night, 150 μmol.m^−2^.s^−1^) for 2-3 weeks before being transferred to a long day growth chamber (16 h light, 21°C, 150 μmol.m^−2^.s^−1^) to induce flowering. Note that seedlings were transferred from short day to constant light for the experiments of Fig. S5C and S8C (24h light, 16°C, 150 μmol.m^−2^.s^−1^). Seeds were staged every-day for up to 10 days by marking the opening of the flower with a cotton thread.

### Measurements of endosperm turgor by microindentation

Siliques were opened and seeds were placed individually on adhesive tape on a microscope slide and covered with water. The slides were then placed on the extended stage of the microindenter (TI 950 Triboindenter, Hysitron). A truncated conical tip with a flat end of ~100 μm diameter (nominal value = 96.96 μm) was used for indentation. The ‘displacement-controlled’ mode was used to allow imposition of a maximum indentation of 30 μm with a load rate of 6 μm/s (5 s extend, 5 s retract). High-resolution force-displacement curves were recorded with a data acquisition rate of 200 points/s. When all of the indentations had been performed, the water was removed and replaced by a drop of clearing solution to allow subsequent embryo staging (see clearing section). Endosperm pressure was calculated using the following formula as described in (*5*):

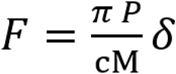

Where F corresponds to the measured force, P to the pressure, δ to the displacement (indentation), and c_M_ to the mean curvature of the presumed load-bearing cell wall.

### Extraction of seed curvature

The membrane marker *Lti6b:GFP* was used to determine the radius of curvature of developing Col-0 and *iku2* seeds by confocal microscopy. Individual seeds were placed on adhesive tape on a microscope slide and covered with water. Confocal imaging was performed on a Leica SP8 upright confocal microscope equipped with a 25x water immersion objective (HCX Fluotar VISIR 25x/0.95 W). The GFP was excited with a LED laser emitting at a wavelength of 488 nm (Leica Microsystems). The signal was collected at 495-555 nm for GFP. For large seeds, several z-stacks were taken and stitched with the LAS (Leica Acquisition System) software. The following scanning settings were used: pinhole size 1AE, 1.25x zoom, 15% laser power, 8000 Hz scanning speed (resonant scanner), frame averaging 4 – 6 times and z intervals of 0.5μm. After imaging, the water was removed and replaced by a drop of clearing solution to allow subsequent embryo staging (See clearing section). The curvature of the seed was extracted using a custom script developed with the ImageJ software. Seed contours were automatically segmented on “by default” thresholded Z-stack projections (Sum-slices) and rotated to align the ellipse-fitting major axis with the Y axis. XZ and YZ orthogonal views at the centroid of the seed were displayed and an ellipse was manually drawn to best fit the surface of the turgid compartment of the seed. The radius of curvature was then calculated using RC = (major axis radius)^2^ /(minor axis radius) for longitudinal and transverse curvatures.

### Quantification of *pELA1::3X-VENUS-N7* expression in ad-oi nuclei

The samples were prepared as described in the previous section. Z stacks of Col-0 and *iku2* seeds expressing *pELA1:3X-VENUS-N7* were acquired using a Leica SP8 upright confocal microscope equipped with a 40x water immersion objective (HCX APO L UV 40x/0.8 W). After imaging, the water was removed and replaced by a drop of clearing solution to allow subsequent embryo staging (See clearing section). Nuclear Fluorescence intensities were measured using a custom-made macro script developed in ImageJ where the nuclei were segmented using on z-stack projections (Sum-slices) using a marker-based watershed (https://imagej.net/Marker-controlled_Watershed).

### Seed clearing, size measurements and embryo staging

To visualize developing seeds, the siliques were opened with a needle, and the seeds were removed with forceps and put in a drop of clearing solution (1 vol glycerol / 7 vol chloral hydrate liquid solution, VWR Chemicals) between a slide and a coverslip. The samples were incubated at least 24h at 4°C before being imaged with a Zeiss Axioimager 2 equipped with a 20x DIC dry objective. The area of the seed was measured by outlining manually the seed using the polygon selection in ImageJ. To compare experimental data with simulations, the radius of the seed was calculated from the measurements of the area by considering the seed as a circle (Area = π (radius)^2^). Seeds were also manually classified based on the developmental stage of their embryo.

### Immunolocalization of cell wall components

Seeds were fixed in ice-cold PEM buffer (50mM PIPES, 5mM EGTA and 5mM MgSO4, pH 6.9) with 4% (w/v) paraformaldehyde. The samples were placed under vacuum (3 x 30 min on ice), rinsed in PEM buffer, dehydrated through an ethanol series and infiltrated with increasing concentrations of LR White resin in absolute ethanol (London Resin Company) over 8 days before being polymerized at 60°C for 24h. The samples were sectioned (1.0 μm thickness) using a diamond knife 45° angle (Diatome, LFG Distribution) mounted on a Leica RM6626 microtome and dried on glass slides.

For JIM5 and LM19 antibodies (Plant Probes), the sections were initially blocked in a PBS solution with 3% (w/v) BSA for 1 hour at room temperature. For 2F4 antibody (Plant Probes), the sections were initially blocked in TCAS buffer (20 mM Tris-HCl, pH 8.2, 0.5 mM CaCl_2_, 150 mM NaCl) with 3% (w/v) skimmed milk for 1 hour at room temperature. The antibodies were applied to the sections overnight at 4°C in a humid chamber. The JIM5 and LM19 antibodies were diluted 1:10 (v/v) in PBS/BSA 1% while the 2F4 antibody was diluted 1:5 in 1% skimmed milk in TCAS buffer. The sections were then washed in an excess of the buffer to dilute the antibody and subsequently incubated for 1 hour at room temperature with the secondary antibody (anti-rat IgG Alexa 488, anti-rat IgM Dylight Alexa 488, and anti-mouse IgG Alexa 488 for JIM5, LM19 and 2F4 respectively) diluted 1:100 in the same buffers as the ones used for diluting the primary antibody. The sections were washed in buffer solutions as described above and finally covered with PBS or TCAS buffer. The samples were then counterstained with filtered Calcofluor White M2R (fluorescent brightener 28; Sigma-Aldrich) at 0.25 mg.ml^−1^ and mounted with VECTASHIELD (Eurobio). The sections were imaged using a Zeiss Axioimager 2 equipped with a 40x dry objective.

The quantification of the immunofluorescence signal was performed with a custom macro script developed on ImageJ software. The two channels were split; the first channel was labelled as the “control” (Calcofluor), and the second channel as the “signal” (antibody) (**Fig.S9A and B**). Testa cells were segmented from the control channel using a stationary wavelet transform and a marker-based watershed (https://imagej.net/Marker-controlled_Watershed). Each testa cell was then manually assigned to its layer (**Fig. S9C**). Enlarged Region Of Interest (ROI) for a given cell (layer n) and for its neighboring cells (layer n-1) were added on a new image and used to define cell wall junctions as being the common region between the two (ImageCalculator (And …) Command). The newly defined ROI was then transferred to the Signal channel for intensity measurements. Finally, cell walls ROIs were overlayed to the composite image of control and signal channels to manually check that the segmentation and localization of the walls had been correctly performed (**Fig. S9D**).

For toluidine blue staining, the sections were incubated for 20 seconds at 70°C with filtered Toluidine Blue 1% / 1% borax before being rinsed with distilled water, dried and mounted in with Entellan mounting medium (Merck). The sections were imaged with a Zeiss Axioimager 2 equipped with a 20x dry objective.

### Generation of the *pIKU2::3X-VENUS-N7* line

The *IKU2* promoter was amplified using the primers Prom-IKU2-B4 (5’-ggggacaactttgtatagaaaagttgGGTCTCTCTTGATAACGATTTG-3’) and Prom-IKU2-B1R (5’-ggggactgcttttttgtacaaacttgTGTTCTCTACGTCGGAAGG- 3’) and cloned into *pDONR-P4-P1R* (Life Technologies). A triple LR Gateway reaction (Life Technologies) was then performed using the *pIKU2-pENTR-R4-L1*, *3X-VENUS-N7-pENTR-L1-L2*, and *3’-ter-pENTR-R2-L3* plasmids as entry vectors and the *pH7m34GW* plasmid as destination vector to generate a *pIKU2::3X-VENUS-N7-pH7m34GW* construct (conferring Hygromycin resistance in plants).

### Genotyping

The *iku2-2* mutant allele was genotyped using the following primers: iku2-Del-For (5’-TTGCTGGAGAAGCTTGTTCTAG-3’) and iku2-Del-Rev (5’-GAACTCCATGGGAATA-TTCCAG-3’).

### Statistical analysis

All experiments were carried out independently at least two times and pooled, unless stated otherwise and each seed corresponds to a biological replicate. Statistical analysis were carried using the R software. In boxplot representations, the midline represents the median of the data while the lower and upper limits of the box represent the first and third quartile respectively. The bars represent the distance between the median and one and a half time the interquartile range. When the number of biological repeats was low (for the pressure, the analysis of *ELA1* expression and the immunolocalizations), individual measurements were superimposed as points on the boxplots. For the remaining representations, points, often connected with lines, correspond to the mean and the error bars to the standard deviation. The relative seed growth rate at day (n) was calculated using the following formular: Relative growth rate = (Area(Day_n_) – Area (Day_n-1_))/ Area (Day_n-1_).

### Data and code availability

All of the experimental data from this article are available upon request to the corresponding author. All the code is available at https://gitlab.inria.fr/mosaic/publications/seed_sup_mat. All the data is available in the main text, in the supplementary materials or at https://zenodo.org/record/4620948#.YFR0Hi1h0UE.

## Supplementary text

## S1 System formalization

### S1.1 Geometrical description

To investigate the antagonist effect of mechanical stress on seed development, we derived the leanest model possible. To that end, we assimilated the seed coat to a spherical shell of radius (*R*) and homogeneous thickness (*H*), yielding a one-dimensional geometrical description of the seed (Fig. 1.C).

### S1.2 Mechanical assumptions

From a mechanical perspective, we assumed a homogeneous and isotropic elastic response of the seed coat to external loading. This enabled us to account for the seed coat overall elastic properties through a single parameter: its effective bulk rigidity modulus (*K*). Assuming furthermore the linearity of this elastic response yielded the well known Hooke law, equation (SE1), relating strain (*ε*) and stress (*σ*) to the effective bulk rigidity modulus within the seed coat:

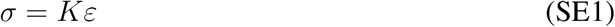

The endosperm influence on the seed coat is limited to an hydrostatic pressure (*P*) applied to the shell from within.

We also considered growth to be a *quasi-static* phenomenon, *i.e.* a continuous succession of mechanical equilibria where the elastic response of the seed coat balances the pressure forces generated onto it by the endosperm.

Given the spherical symmetry of our representation, this assumption yields the Laplace law, equation (SE2), relating tensile stresses within the seed coat to the endosperm pressure and the geometrical properties of the shell:

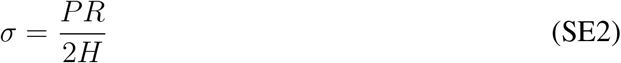

### S1.3 Biological assumptions

We considered two responses of the seed coat to the mechanical solicitation: Growth and cell wall stiffening.

Both mechanisms are complex, high-level, processes involving numerous biomechanical and biochemical entities and mechanisms; *e.g.* cell wall polymers and enzymes, cytoskeleton assemblies, transcription factors, transmembrane carriers. A formalization accounting exhaustively for all the molecular processes at stake is of course out of reach and not in the scope of this work. To alleviate this complexity we opted for an *empirical* formalization of both phenomena.

#### Cell growth

We described cell growth within the seed coat with a thresholded strain-based law, equation (SE3).

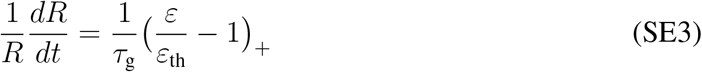

Equation (SE3) states that the relative growth rate of the seed coat is proportional to the strain above a given threshold (*ε*_th_). If the strain remains below this threshold, the growth rate vanishes. The growth characteristic time (*τ*_g_) quantifies the kinetics of this irresversible expansion. This empirical law can be seen as an extension of the orignal Lockhart (*6*) and Ortega (*7*) models. It has been originally developed for 3D Finite Element models (*8–10*) and accounts well for experimental observations, namely that cells expand orthogonally to the cell wall stiffest direction.

#### Cell wall stiffening

We formalized cell wall stiffening within the seed coat as a first-order ordinary differential equation, equation (SE4), expressing the stiffening rate of the cell wall as the combination of a *production* and a *degradation* term.

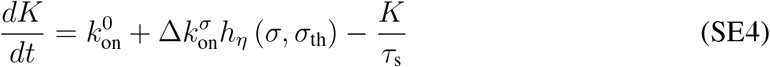

The production term is composed by the first two elements of the right hand-side of equation (SE4). The first one depicts a *basal* stiffening rate. Combined with de degradation term, third element of equation (SE4) *rhs*, it provides the seed coat with a stationary value 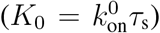 for its effective bulk rigidity modulus; when no mechanobiological regulation is at play.

The second element of equation (SE4) *rhs* accounts for the the seed coat stress-sensitive stiffening ability. It corresponds to a *Hill function*, equation (SE5), increasing the production rate from 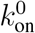 to 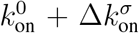 when tensile stresses within the seed coat go from low values (*σ ≪ σ*_th_) to high ones (*σ ≫ σ*_th_).

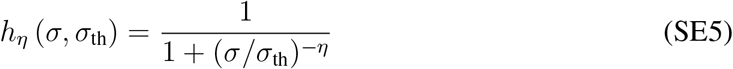

The parameter *σ*_th_ can be interpreted as a threshold for this stress-sensitive mechanism and the Hill exponent *η* as a measure of its non-linearity (*i.e.* the bigger *η* the sharper the response to stress).

All parameters and variables used through equations (SE1) to (SE5) are listed within table (ST1).

### S1.4 Dimensionless formalization

To analyze the properties of the differential system composed by equations (SE3) and (SE4) we derived a dimensionless version from it.

By combining equations (SE1) to (SE4) and normalizing the three variables (*t, R, K*) respectively by (*τ*_g_*, H, K*_0_) we get the desired dimensionless description grasped through system (SE6).

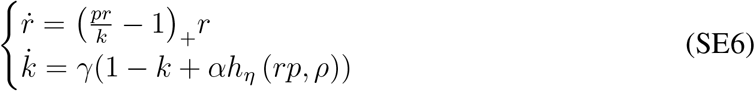

#### N.B.

In system (SE6) derivation with respect to the dimensionless time variable 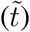 is noted with a dot over the derived quantity.

All the dimensionless parameters and variables of system (SE6) are listed within table (ST2).

#### Rationale

This dimensionless approach seems relevant in our case, for several reasons: It simplifies the system by removing intermediate variables (*σ* and *ε*) and it highlights the relationship between the structural properties of the equations and the dynamical properties of the system. But most importantly, since Equations (SE3) & (SE4) derive from empirical considerations, their parameters cannot be tracked to actual biochemical and/or rheological properties that could be properly measured experimentally. Parametrizing equations (SE3) & (SE4) with relevant values appears therefore difficult. By focusing on dimensionless version of these equations, we alleviate this difficulty: The parameters we need to estimate correspond now to ratios between comparable quantities. The drawback is that we can only extract qualitative information from their analysis. For instance, the condition: *γ* = 10 ⇒ *τ*_g_ = 10*τ*_s_, can only be translated into the following qualitative statement: *growth is slower than the stiffening process by one order of magnitude*.

## S2 Numerical simulations

Together with a set of initial values {*r*_0_*, k*_0_}, equation (SE6) constitutes an *Initial Value Problem*. Given a set of values for the parameters {*α, γ, ρ, η*} and the control variable *p*, one can simulate the seed growth dynamics by resolving such *ivp*.

To that end, we implemented system (SE6) in python (v 3.7.5) and make use of the `solve_ivp` method from the `scipy.integrate` module (v 1.3.1) to solve it given some initial conditions and pressure value. All simulations were performed between *t*_min_ = 0 and *t*_max_ = 3 with a time step *δt* = 0.02.

### Matching simulation time with experimental time

The simulation time is measured in units of growth characteristic time (*τ*_g_). This notion is rather arbitrary and does not need to be specified to perform simulations (only the ratio between growth and stiffening characteristic times, namely parameter *γ* is needed). However, in order to compare simulation results with experimental measurements we set a *convertion factor* in order to express simulation times in *DAP/DPA* unit:

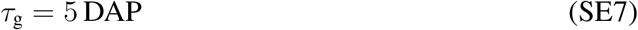

### Code availability

All simulation scripts as well as data analysis notebooks are freely available on line: https://gitlab.inria.fr/mosaic/publications/seed_sup_mat. A detailed description of how to install and run our simulations is provided within the `README.md` file within this repository.

### Data availability

Simulation scripts and notebooks require input data. These are available on line as well: https://zenodo.org/record/4620948#.YFR0Hi1h0UE

### S2.1 Parameter space exploration

As mentionned above, we started this study with no clear assumptions concerning the values of the four parameters featured in the stiffening equation, second line of system (SE6).

We therefore investigated system (SE6) behavior for a wide range of parameter values, see table (ST3). Overall, we sampled our four dimensional parameter space into 5 · 10^5^ parameter sets and simulated the dynamics of the system for all of them. All simulations, within this parameter space exploration, featured the same, constant value of relative pressure: *p* = 1.2, chosen arbitrary, slightly above the threshold value 1.

To that end, we used the python library `pypet` (*11*) to distribute and manage simulations over a multi-core computing server. Data management was performed using the `DataFrame` data structure from the `Pandas` library (v 0.25.3) (*12*). Data processing and analysis were performed within `Jupyter notebooks` (v 6.0.2) and visualization with the `seaborn` library (v 0.9.0) (*13*).

#### Selection of the parameter space region to investigate

Due to computational limitations, we had to limit the range of our parameter space exploration.

- the *α* parameter quantifies the amplitude of the *stress-sensitive* stiffening term, compared to the *passive* terms, in the second line of system (SE6). Theoretically, the only constrain on its value is that it belongs to ℝ^+^. But AFM measurements performed during organogenesis at the Shoot Apical Meristem, reported 3 to 4 fold variations of wall stiffness value (*14*), suggesting that the stress-sensitive term should be significant but not overwhelming. To that end, we tested values between 1 and 11.
- We considered odd integer Hill function exponents (*η*) ranging from 3 to 9 to probe the influence of the non-linearity of the stress-sensitive stiffening term.
- The parameters *γ* and *ρ* correspond respectively to the ratio of the characteristic times and threholds between the growth and stiffening processes. We chose to sample them along a logarithmic scale, *i.e.* by setting *γ* = 10^*x*^ and *ρ* = 10^*y*^ and considering ranges centered on 0 for *x* and *y*. Precisely, we chose: *x, y* ∈ [−2.5, 2.5]. This enabled us to consider symmetric situations with respect to the kinetics of stiffening compared to growth.

For the three parameters *α*, *ρ* and *γ* we sampled the considered intervals into 50 points each. Combined with the four considered values for the parameter *η*, the four dimensional region we considered within the parameter space has been discretized into 5 *·* 10^5^ samples, each corresponding to a unique set of values {*α, η, ρ, γ*}.

#### Result filtering

Once this systematic exploration done, we discarded simulations that did not meet the two following criteria:

- Simulations must have converged toward a steady value, first line of equation (SE8).
- The radius final value must lie within a range compatible with experimental measurements, second line of equation (SE8).

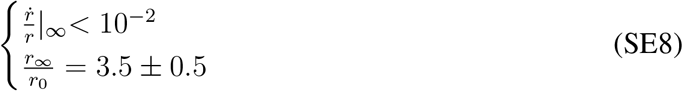

Once this filtering done, less than 2 · 10^3^ simulations remained. These simulation results and parameter values are stored within the *sim res cstePressure highResParamScan.hdf5* `DataFrame` within the `/model/data/results/` folder accessible on the gitlab repository associated with this manuscript.

#### Comparison with experimental data

We then compared the relative radius dynamics of each kept simulation to experimental measurements. To that end, we first matched the simulation time with the experimental one by applying the following change of variable: 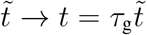 (with 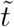 represents the simulation time), with the value for *τ*_g_ given in expression (SE7). Then, we sampled every simulations at integer time steps (corresponding to experimental sampling times): *t_k_* ∈ {0, 1, 2*, …* 10} and then measured the (root-mean-square) distance between the following vectors:

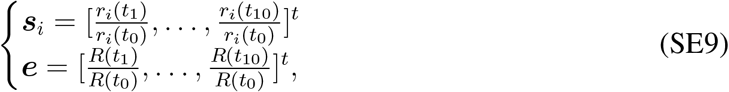

where the index *i* runs over all kept simulations and *R*(*t_k_*) depicts the mean value of the seed radii measured at time step *t_k_*.

#### N.B.

*One can note that the simulation result vector **s**_i_ is constructed from the relative radius variable r = R/H, while the experimental measurement vector **e** is directly constructed from the seed radius estimation R. In order to compare them, we assumed seed coat thickness (H) constant during the seed expansion phase. This assumption was mainly motivated by the fact that the number of cell layers within the seed coat remains constant and that each cell layer, within the seed coat, roughly keeps the same thickness during the studied period.*

We then defined a fitting score (*F* (*i*)) for each simulation (*i*) as the inverse of the distance between the corresponding vector ***s**_i_* and the vector of experimental measurements ***e***:

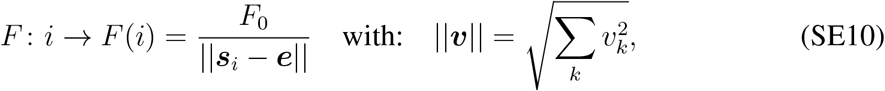

where the constant *F*_0_ = min(∥***s**_i_* − ***e***∥) is used in order to normalize the fitting score to one for the best fitting simulation.

This fitting procedure enabled us:

- To visualize sub-regions of the parameter space corresponding to simulations matching the dynamical properties of actual seeds, see figure (1.D) within the main text and supplementary figure (S4).
- To sort all of the kept simulations by their degree of similarity to experimental data. And concentrate our analysis of parameter values on the hundred best-fitting simulations, see figures (1.B) and (3.A) within the main text.

We performed this fitting analysis against three sets of experimental measurements, corresponding to three different genotypes: wild type (ecotype Col-0), *iku2* and *ede1-3*. The results are given in table (ST4).

### S2.2 Simulations with time dependent pressure

#### Rationale

While the assumption of constant endosperm pressure appears relevant to model the *iku2* mutant, experimental data suggest that endosperm pressure is a monotonously decreasing function of time in the *WT* case. Since *iku2* expression is restricted to the endosperm compartment, Fig. (S5.A), the discrepancy between the *WT* and the *iku2* phenotypes cannot be accounted for through different values for the set of parameters ({*α, η, ρ, γ*}) within our model, for these parameters solely grasp properties of the seed coat.

Based on these facts, we wonder if we could recover the WT growth behavior from the *iku2* best-fitting parameters values but combined with a time-decreasing pressure function instead of a constant pressure value.

#### Results

The purple graph in (Fig.2E) shows the pressure drop function we implemented, compared to the constant value initially used to match the *iku2* data. It consists of a 30% drop from the initial value, spanned between 1 and 5 *DAP*. Such a drop qualitatively corresponds to the pressure variation observed between *iku2* mutants and WT seeds in (Fig.2C).

As (Fig.3B) shows, replacing the constant pressure with a time-decreasing function enabled us to increase the final radius within our simulations in a way comparable to the actual dynamics of WT seeds.

## Tables

**Table ST1:** Variables & parameters.

| Param. & Var. | Meaning | Units | Values<br>Ref. |
| --- | --- | --- | --- |
| **Geometry** |  |  |  |
| $R$ | Radius of the growing spherical shell | $\mu\text{m}$ | 47.5 – 310 (measures) |
| $H$ | Thickness of the shell | $\mu\text{m}$ | 20 – 25 (measures) |
| **Mechanics** |  |  |  |
| $P$ | Pressure exerted by the endosperm on the seed coat | MPa | $(3 - 18) \cdot 10^{-2}$ (measures) |
| $K$ | Young's modulus of the shell | MPa | $N/A$ |
| **Growth** |  |  |  |
| $\tau_g$ | Growth characteristic time | s | $N/A$ |
| $\varepsilon_{th}$ | Growth threshold (strain) | % | $N/A$ |
| **Stiffening** |  |  |  |
| $\tau_s$ | Cell wall stiffening characteristic time | s | $N/A$ |
| $\sigma_{th}$ | Stiffening threshold (stress) | MPa | $N/A$ |
| $k_{on}^0$ | Intrinsic stiffening rate | $\text{MPa} \cdot \text{s}^{-1}$ | $N/A$ |
| $\Delta k_{on}^\sigma$ | mechanosensitive increase of the stiffening rate | $\text{MPa} \cdot \text{s}^{-1}$ | $N/A$ |
| $\eta$ | Hill function exponent, steepness of the mechanosensitive stiffening mechanics | ∅ | $N/A$ |
| $K_0$ | Stationnary value of the bulk rigidity modulus without mechanosensitive stiffening at play. | MPa | $N/A$ |

**Table ST2:** Dimensionless variables & parameters used within the dimensionless system, see SE6. The given values and ranges correspond to the ones used and explored within the parameter space simulations.

Table ST1: Variables & parameters.
|  | Name | Symbol | Expression | Interpretation | Value/Range |
| --- | --- | --- | --- | --- | --- |
| **Variables** | Relative time | $\tilde{t}$ | $t/\tau_g$ | Time normalized by the growth characteristic time. | $[0, 3]$ |
| | Relative radius | $r$ | $R/H$ | Radius of the seed normalized by the thickness of the seed coat. | 1.5 |
| | Relative stiffness | $k$ | $K/K_0$ | Seed coat effective stiffness normalized by its stationary value when on stress-induced process is active. | 1 |
| | Relative pressure | $p$ | $P/(2\varepsilon_{th}K_0)$ | Endosperm pressure divided by the growth threshold times the basal value of the effective seed coat stiffness. | $[0.5, 2]$ |
| **Parameters** | Stress-stiffening strength | $\alpha$ | $\Delta k_{on}^\sigma/k_{on}^0$ | Maximum relative stiffness increase due to the stress-induced mechanism. | $[1, 11]$ |
| | Characteristic time ratio | $\gamma$ | $\tau_g/\tau_s$ | Ratio between growth and stiffening characteristic times. | $[10^{-2.5}, 10^{2.5}]$ |
| | Threshold ratio | $\rho$ | $\sigma_{th}/(\varepsilon_{th}K_0)$ | Ratio between the stiffening and growth thresholds. | $[10^{-2.5}, 10^{2.5}]$ |
| | Hill exponent | $\eta$ | — | Steepness of the mechanosensitive stiffening mechanics. | $\{3, 5, 7, 9\}$ |

**Table ST3:** Details of the sampling used to perform the parameter space exploration.

| **Variable** | **Min. value** | **Max. Value** | **Increment** | **Samples** | **Scale** |
| --- | --- | --- | --- | --- | --- |
| $\alpha$ | 1 | 11 | .2 | 50 | linear |
| $\eta$ | 3 | 9 | 1 | 4 | linear |
| $\gamma$ | -2.5 | 2.5 | .1 | 50 | logarithmic |
| $\rho$ | -2.5 | 2.5 | .1 | 50 | logarithmic |

**Table ST4:** Results from the parameter space exploration campaign: parameter values that correspond to simulations fitting experimental data the best. The mean, median and standard deviation (st. dev.) values have been computed over the 100 best fitting results.

Table ST3: **Details of the sampling used to perform the parameter space exploration.**
|  |  | **WT (Col-0)** | ***iku2*** | ***ede1-3*** |
| --- | --- | --- | --- | --- |
| **Best fit** | **Distance** | 0.26 | 0.18 | 0.33 |
|  | **Simulation Id** | 228781 | 98680 | 186431 |
| **Parameters** | **values** |  |  |  |
| $\alpha$ | best: | 9.20 | 8.80 | 5.80 |
|  | mean: | 7.74 | 7.40 | 7.94 |
|  | median: | 7.80 | 6.30 | 8.50 |
|  | st. dev.: | 1.10 | 1.85 | 1.12 |
| $\eta$ | best: | 7 | 9 | 7 |
|  | mean: | 6.40 | 7.20 | 5.40 |
|  | median: | 6.00 | 7.00 | 5.00 |
|  | st. dev.: | 1.56 | 1.08 | 0.80 |
| $\gamma$ | best: | 1.00 | 0.63 | 2.00 |
|  | mean: | 4.57 | 84.55 | 12.95 |
|  | median: | 2.25 | 53.97 | 14.22 |
|  | st. dev.: | 4.21 | 90.81 | 7.70 |
| $\rho$ | best: | 3.98 | 3.16 | 3.98 |
|  | mean: | 4.89 | 4.27 | 5.71 |
|  | median: | 4.50 | 5.01 | 6.31 |
|  | st. dev.: | 1.01 | 0.91 | 0.95 |

